# Intracortical neural activity distal to seizure-onset-areas predicts human focal seizures

**DOI:** 10.1101/529966

**Authors:** Timothée Proix, Mehdi Aghagolzadeh, Joseph R Madsen, Rees Cosgrove, Emad Eskandar, Leigh R. Hochberg, Sydney S. Cash, Wilson Truccolo

## Abstract

The apparent unpredictability of epileptic seizures has a major impact in the quality of life of people with pharmacologically resistant seizures. Here, we present initial results and a proof-of-concept of how focal seizures can be predicted early in advance based on intracortical signals recorded from small neocortical patches away from identified seizure onset areas. We show that machine learning algorithms can discriminate between interictal and preictal periods based on multiunit activity (i.e. thresholded action potential counts) and multi-frequency band local field potentials recorded via 4 X 4 mm^2^ microelectrode arrays. Microelectrode arrays were implanted in 5 patients undergoing neuromonitoring for resective surgery. Post-implant analysis revealed arrays were outside the seizure onset areas. Preictal periods were defined as the 1-hour period leading to a seizure. A 5-minute gap between the preictal period and the putative seizure onset was enforced to account for potential errors in the determination of actual seizure onset times. We used extreme gradient boosting and long short-term memory networks for prediction. Prediction accuracy based on the area under the receiver operating characteristic curves reached 90% for at least one feature type in each patient. Importantly, successful prediction could be achieved based exclusively on multiunit activity. This result indicates that preictal activity in the recorded neocortical patches involved not only subthreshold postsynaptic potentials, perhaps driven by the distal seizure onset areas, but also neuronal spiking in distal recurrent neocortical networks. Beyond the commonly identified seizure onset areas, our findings point to the engagement of large-scale neuronal networks in the neural dynamics building up toward a seizure. Our initial results obtained on currently available human intracortical recordings warrant new studies on larger datasets, and open new perspectives for seizure prediction and control by emphasizing the contribution of multiscale neural signals in large-scale neuronal networks.

## Introduction

A third of patients with epilepsy are not responsive to medication and experience recurrent seizures through their lives [1]. This number has not changed much in the past decades despite the efforts toward the development of new antiepileptic drugs [2]. The alternative of regional surgical resection to remove identified epileptogenic areas in the brain is only applicable to about 25% of these cases, carry substantial risks, and has limited efficacy [3]. More recently, various studies and clinical trials have started examining new therapeutic approaches based on seizure prediction or early detection for warning systems and seizure prevention and control via closed-loop electrical stimulation [4–6]. These new approaches have relied on intracranial grids covering the cortex over several centimeters and/or depth electrodes. Despite promising initial results [4,5,7], efficacy remains limited.

One of the current barriers to more efficient seizure prediction and control is the lack of understanding of the multiscale neural dynamics and neurophysiological processes leading to seizure events, especially the activity at the level of neuronal action potentials [8–11]. In addition, most previous studies have focused on mechanisms in the seizure onset areas (SOAs) and epileptogenic zones. An overlooked aspect of seizure initiation and maintenance has been the broader engagement and contribution of large-scale neuronal networks beyond these identified areas [12–14].

Here, we address the contribution to seizure prediction of neuronal activity recorded from small neocortical areas away from the identified SOAs. We tackled this problem by evaluating if seizures can be predicted from intracortical neural signals recorded via microelectrode arrays (MEAs) implanted in patients with drug resistant focal seizures undergoing neuromonitoring for resective surgery. We note that, currently, intracortical MEA recordings of human epileptic seizures have been limited to recordings done in patients during the neuromonitoring period preceding the resective neurosurgery, which typically corresponds to one to two weeks approximately. No longer datasets with MEAs are currently available worldwide for epileptic patients. We used machine learning algorithms, specifically long short-term memories (LSTM), a type of recurrent neural network, and extreme gradient boosting (XGBoost), for seizure prediction. We show that seizures can be successfully predicted based on local neuronal population spiking reflected in multiunit activity (MUA), i.e. counts of thresholded but unsorted action potentials from recorded neurons. In addition, local field potentials (LFPs) were particularly predictive in the 0-8 Hz and 50-500 Hz frequency bands, with the latter possibly including MUA features. Notably, different sites in the MEA showed higher predictive power than others according to given features, indicating the existent of a finer spatial microdomain structure in the seizure predictive activity. To our knowledge, this is the first study to systematically demonstrate seizure prediction based on neural activity recorded intracortically and distal to identified SOAs.

## Materials and Methods

### Patients and data acquisition

Research protocols were approved by local Institutional Review Boards at Massachusetts General and at Brigham and Women’s Hospitals (Partners Human Research Committee) and at the Rhode Island Hospital. Informed consent was obtained from all of the patients participating in this study and experimental methods were carried out in accordance with the regulations of the local Institutional Review Boards. Patients were undergoing neuromonitoring for identification of target areas for resective surgery. The clinical aspects and recording setup for the five patients (P1-P5) referred in this study are provided in detail in Wagner et al. (2015). Here, we briefly review this information in Table 1. Typical clinical intracranial EEG recordings, including electrocorticography (ECoG) grids and/or depth electrodes, were performed as part of the neuromonitoring prior to the resective surgeries. Research MEA recordings were obtained from a 4×4 mm^2^ 10 X 10 microelectrode array with 96 recording microelectrodes (Blackrock Inc, Salt Lake City, UT; Fig 1). Electrical potentials were recorded broadband (0.3Hz – 7.5Hz) and sampled at 30 kilosamples/s. Seizure onset areas and times were determined by the clinical teams at Massachusetts General, Brigham and Women's, and Rhode Island Hospitals, independently of the research being reported here.

**Table 1:** Patient information.

| Patient | Gender | Age | Epilepsy type | Symptoms | Side | MRI | Histopathology | Surgical resection | Surgical outcome | Location of the MEA | Number of selected seizures |
| --- | --- | --- | --- | --- | --- | --- | --- | --- | --- | --- | --- |
| **P1** | Male | 45 | Mesial temporal sclerosis | loss of consciousness, motor manifestations | Right | ? | ? | Right temporal lobectomy | I / 4 years | MTG; not in SOA; not in identified epileptogenic zone, but in irritative zone. | 2 |
| **P2** | Male | 32 | Mesial temporal sclerosis | loss of consciousness, motor and autonomic manifestations | Left | Polymicro-gyria | Hippocampal sclerosis | Left temporal lobectomy | I / 3.5 years | STG; not in SOA; not in, epileptogenic zone or irritative zone. | 2 |
| **P3** | Female | 25 | Cortical dysplasia | loss of awareness, autonomic manifestations | Right | Cortical dysplasia | Focal superficial cortical gliosis, areas of encephalomalacia | Extensive right temporal resection | I / 3 years | STG; not in SOA; not in epileptogenic zone, but unclear if in irritative zone. | 3 |
| **P4** | Male | 21 | Mesial temporal sclerosis | loss of awareness, motor components | Left | Normal | Gliosis, and moderate neuronal loss | Left temporal lobectomy | I / 4 years | STG; not in SOA; not in epileptogenic zone or in irritative zone. | 2 |
| **P5** | Female | 52 | Mesial temporal sclerosis | loss of consciousness, motor components | Left | Encephalomalacia | Hippocampal sclerosis | Resection of left anterior lobe | I, then resumed at lower frequency | MTG; not in SOA; not in epileptogenic zone, but in irritative zone also showing sclerosis (no dysplasia) in superficial cortical layers. | 2 |

MTG/STG: middle/superior temporal gyrus; SOA: seizure onset area; The last column refers to the number of seizures included in this study based on the adopted constraints (e.g. time period between seizures, etc. See Materials and Methods for details).

**Fig 1.**
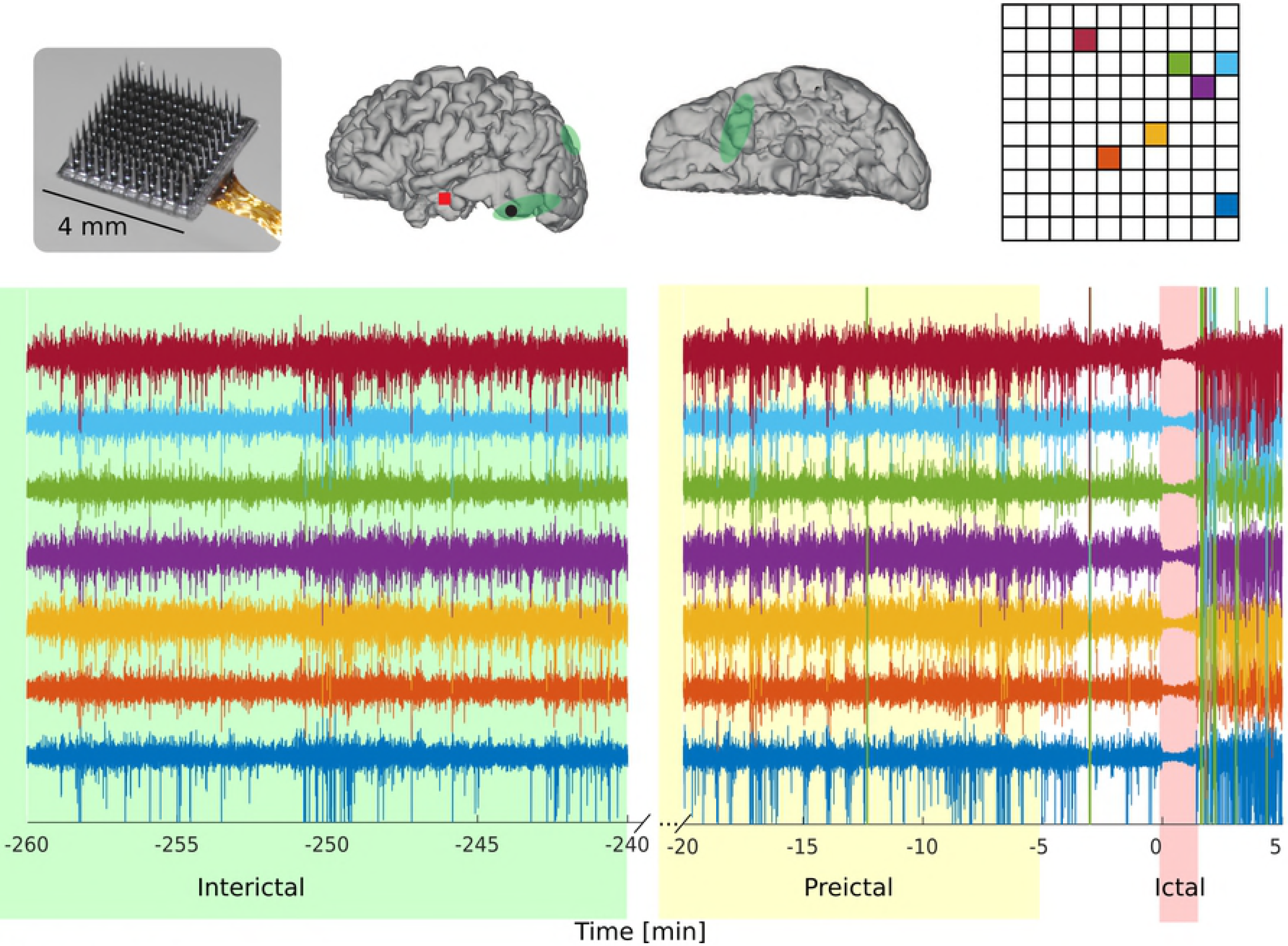
MEA recordings prior and during a seizure. A 4 X 4 mm2 MEA (top left) was intracortically implanted in the middle gyrus of the temporal lobe in patient P5 (middle panel; the red square denotes the location of the MEA on the reconstructed cortical surface). LFP Field potential time series are shown for selected MEA channels prior and during seizure (bottom, colors corresponding to the microelectrode sites shown on the top right). The interictal period is chosen four hours away from any seizure and the preictal period is defined as the 1-hour period between 65 and 5 minutes prior to seizure onset.

### Preprocessing

Broadband MEA recordings were preprocessed offline to obtain three main types of signals: MUA counts, MUA envelope and LFPs (Fig 2). MUA counts for each recording electrode site were obtained by first high-pass filtering (> 250 Hz) the broadband recordings and then counting threshold crossings (- 3 SD) in Δ=0.5 ms time bins. To obtain the MUA envelope signal we followed the general procedure described in [15] with minor adaptations. The high-pass filtered signal was clipped at ±3 SD to attenuate the contribution of large amplitude action potentials from neurons very close to the microelectrode tip and emphasize the broader population activation. The signal was then rectified (squared) and low-pass filtered at 10 Hz. LFPs were obtained by low-pass filtering (<500 Hz) the broadband recordings. All filtering operations were performed with a Butterworth filter (9^th^ order; zero phase design to improve stability; Matlab, Mathworks). LFPs, MUA count (0.5 ms time bins) and MUA envelope were sampled at 2 kilosamples/s.

**Fig 2.**
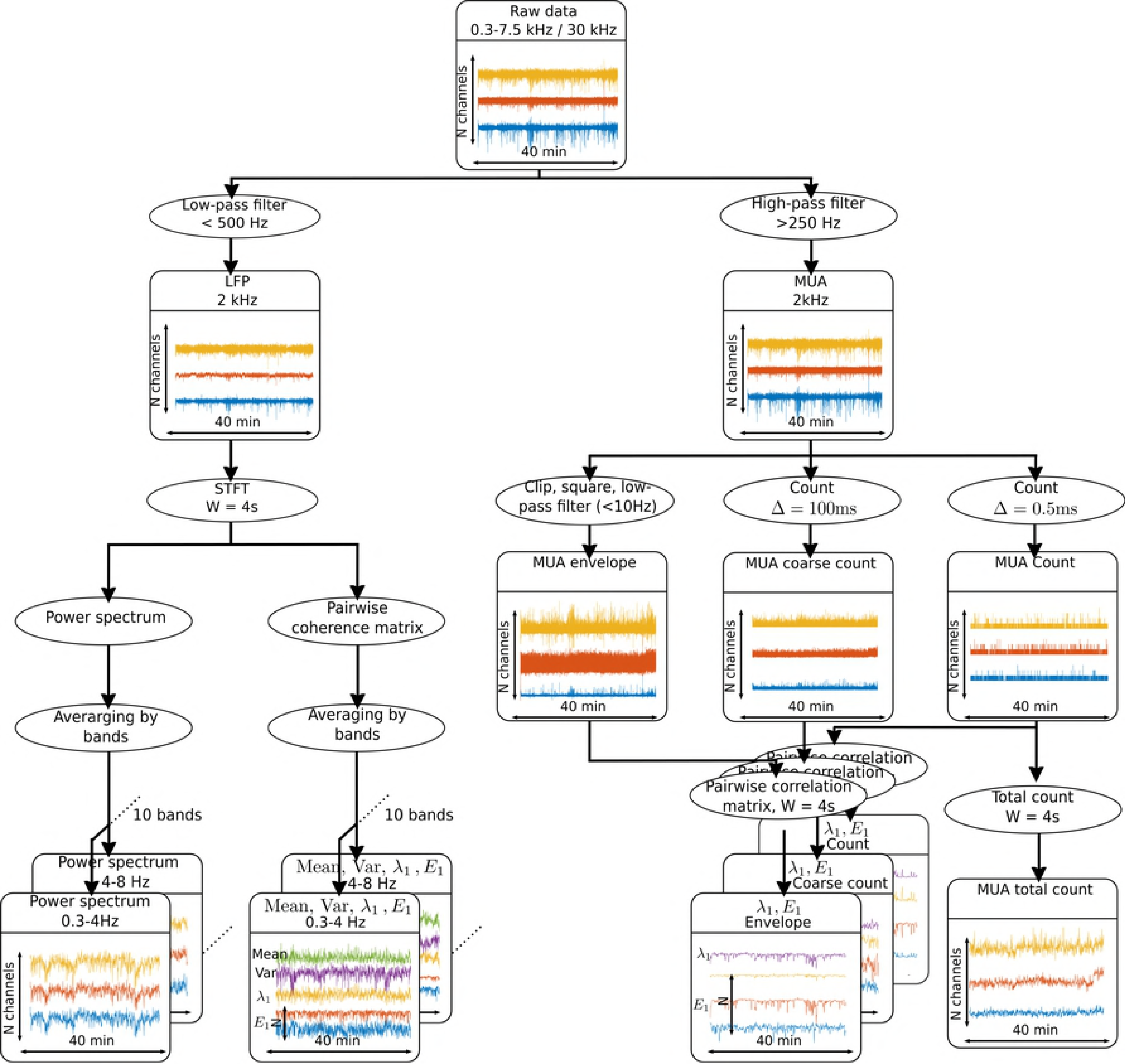
Processing pipeline for extracting features from the MEA recorded signals. The broadband field potential signal is low-pass and high-pass filtered to obtain the LFP and the MUA, respectively. For LFP (left), a short-term Fourier transform (STFT) was applied to obtain the power spectrum, as well as the mean, variance, and leading eigenvalues and eigenvectors of the pairwise spectral coherence matrix averaged over frequency for each frequency band. For MUA (right), the count for each channel, the total count across channels, as well as the leading eigenvalues and eigenvectors of the pairwise correlation matrices for the MUA envelope, the coarse MUA count and the MUA count were computed.

### Feature extraction

Features were extracted from the MUA count, MUA envelope and LFP channels (Fig 2). LFP power spectrum and pairwise spectral coherence were computed via multitaper methods [16] in consecutive W=4 second time windows (2-s overlap) and a half-bandwidth of 2Hz. Four groups of features were extracted: (i) the power spectrum of the LFP averaged across different frequencies in each of ten defined frequency bands (0.3-4 Hz, 4-8 Hz, 8-12 Hz, 12-18 Hz, 18-25 Hz, 25-50 Hz, 50-80 Hz, 80-150 Hz, 150-300 Hz, 300-500 Hz), henceforth called LFP power spectrum features; (ii) the leading eigenvalue and eigenvector of the pairwise coherence matrix for each frequency band (coherence values were averaged across different frequencies in the same band), as well as the mean and variance coherence values computed over the coherence matrix for a given frequency band (henceforth called LFP coherence); (iii) the total number of MUA counts (henceforth referred to as the total MUA count) in each W = 4 second time window for each channel; (iv) the leading eigenvalue and eigenvector of the pairwise correlation matrices for MUA count, MUA coarse count, and MUA envelope. The MUA coarse count was obtained by computing the spike counts in coarser time bins of Δ=100 ms. Pearson correlation coefficients were computed based on the 4-second count series for each channel pair. Correlations were computed over a range of time lags, up to ±50 ms. For each channel pair the extremum of the lagged correlation function was selected for each time window. We refer to those features (leading eigenvalue/eigenvector of correlation matrices) as MUA correlation features. These sets of features allowed us to assess the contribution of features directly related to activation patterns across the MEA (i.e. LFP power spectrum and MUA count) and of features related to pairwise co-activation patterns (second order statistics) across the MEA (i.e. LFP spectral coherence and MUA correlation).

### Datasets and leave-one-seizure-out cross-validation

As stated above, we formulate the seizure prediction problem in terms of classifying selected features into interictal and preictal. Prediction moved forward at 2-second time steps. Consistently with two previous seizure prediction studies [7,17], the labeled interictal time-blocks were restricted to be at least four hours away from any seizures in order to attenuate potential time overlap between interictal and preictal or interictal and postictal states. Also following [7,17], preictal segments were defined as the one-hour interval between 65 and 5 minutes prior to seizure onset (Fig. 1).

We used a leave-one-seizure-out cross-validation procedure. Specifically, one set of interictal and preictal data segments corresponding to a particular target seizure were left out of the training set and used for testing. The same procedure was then iterated for each of the seizures from each patient.

### Classification using LSTMs

We used a small LSTM network [18,19] followed by a dense layer with two units for classification. The equations for a single LSTM cell are (schematics in Supplementary Fig 1):

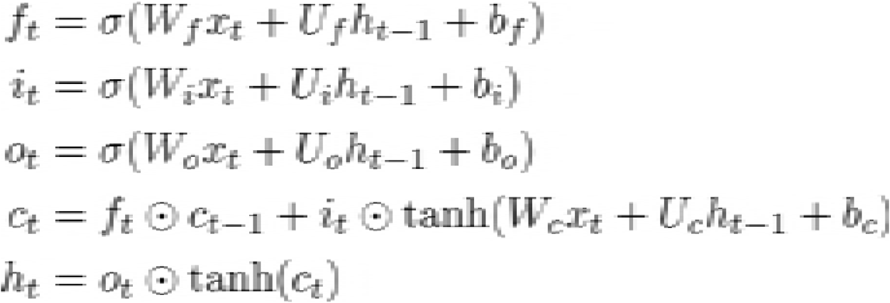

where ⊙ denotes the entrywise (Hadamard) product, with *x*_*t*_, *h*_*t*_, *c*_*t*_, corresponding to the cell’s input, output and state, respectively; *f*_*t*_, *i*_*t*_ and *o*_*t*_, correspond to the forget, input and activation gates, respectively; *W*_*v*_, *U*_*v*_, and *b*_*v*_ are learned parameters for *v* ∈ {*f,o,i*}. The terms *σ(.)* and *tanh(.)* denote the sigmoid and hyperbolic tangent output functions, respectively. Overfitting was controlled by adopting the “dropout” parameter approach commonly used in deep learning. Specifically, the dropout rate for both *W* and *U* were set to 20% during training. In addition, we reduced overfitting by choosing a small network of only 3 cells. In this way, the number of parameters of the LSTM network plus the dense layer, given by 4*((#features+1).#cells + #cells^2^) + 2#cells + 2*, remained smaller than the number of training samples. The number of training samples ranged from 6.938 to 90.138, while the corresponding number of parameters varied from 1.208 to 11.576. In addition, the small network size also facilitated visualizing the activity of the LSTM cell states over time. We used an “Adadelta” optimizer and cost-sensitive learning with class weights corresponding to the class imbalance between positive and negative examples in the training set. We used the LSTM algorithm as implemented in Keras 1.2.1. Throughout this study, we varied only one hyperparameter, which corresponded to the length of the history of the input to the network (or equivalently the number of time iterations for the LSTM network). We explored a range of values including 2, 3, 5, 10, 15, or 25 consecutive four seconds time windows with 50% overlap, which resulted in the history extending from 6 to 52 seconds in the past.

To prepare training and testing datasets for the LSTM network fitting and performance assessment, the feature segments must be sliced in blocks of consecutive time points, whose length is given by the length of the history of the input to the LSTM network. Seizures are typically rare events, resulting in very imbalanced datasets, i.e. many more interictal than preictal samples. We attenuated this imbalance issue by oversampling the preictal class, specifically by overlapping between 50% and 96% the data segments (time windows) labeled as preictal in the training dataset depending of the size of the history used in the LSTMs. Nevertheless, no overlapping between segments was applied to preictal test data for the assessment of predictive power.

### Assessment of classifiers’ seizure-prediction performance

To assess the performance of the classifiers, we computed the area under the receiver operating characteristic (ROC) curve, henceforth referred to as AUC, which evaluates the true positive and false positive rates for varying thresholds.

The performance of a chance-level predictor was obtained by Monte Carlo random permutations as follows. Since the different samples are not statistically independent, we cannot randomly shuffle all the labels of the training samples fed to the LSTM without losing correlations between samples. To mitigate this effect, we divided the training set in non-overlapping 10-minute segments, and randomly shuffled the labels for these 10-minute segments instead. These segments were then processed as described above, and used to train the LSTM network. This operation was repeated N = 200 times to obtain an AUC distribution under the null hypothesis of chance level prediction. Here, we report the corresponding p-values and statistical significance for each patient and group of features separately; these p-values were computed according to *p = (1 + #{AUC*>AUC}) / (1 + N)*, where AUC was computed on the true test dataset and the {AUC*} set was computed based on the chance-level surrogate datasets.

We applied the false discovery rate (FDR) to correct for the multiple testing [20], with chosen targets of *α=0.05* and *α=0.01.*

### XGBoost feature space reduction

To evaluate the contribution and predictive power of different features, we used the XGBoost algorithm [21] on the same training and testing datasets as for the predictions based on the LSTM networks. XGBoost parameters were set to default values, except for the maximum depth of a tree (“max_depth=3”), and for the objective function which was set to the logistic function (“objective=binary:logistic”). A cost-sensitive learning with class weights corresponding to the class imbalance between positive and negative examples in the training set was used as well (“scale_pos_weight=class_imbalance_ratio”). The same time series blocking scheme as for the LSTM network was used for the XGBoost algorithm. Note that time consecutive features were considered as time independent features in the XGBoost algorithm. Feature importance for both the LFP power spectrum and LFP pairwise spectral coherence was evaluated by examining different frequency bands separately and by summing up and normalizing all the spatial components (across the MEA) of the features for each frequency band. Feature importance was assessed for each group of features by summing up and normalizing all the obtained feature importance for each spatial component. We used the XGBoost algorithm as implemented in XGBoost 5.2.0.

### LSTM models based on a reduced set of features according to XGboost ranking of feature importance

We also examined how a reduced set of features, selected from the most important features obtained by the XGBoost feature space reduction, affected the performance of LSTM prediction. This reduced feature set was then used in the analyses reported in the Results section. The selected subset included: LFP power spectrum (bands 50-80 Hz, and 80-150 Hz), leading eigenvectors from the LFP pairwise spectral coherence matrices (bands 0-4 Hz, and 300-500 Hz), MUA total count, and the leading eigenvectors from pairwise correlation for the MUA envelope.

## Results

### Local LFP and MUA predict seizures away from the seizures onset

We first evaluated if seizures could be predicted from local neuronal activity away from the determined SOAs. Broad-band field potentials were recorded from 10 X 10 microelectrode arrays (MEAs, 4 X 4 mm^2^) implanted in five patients with epilepsy (Fig 1 and Table 1). In all cases, the MEA location was two to three centimeters away from the SOAs identified by the clinical team.

For each recorded clinical seizure, the preictal period was defined as the interval between 65 and 5 minutes prior to the seizure onset, and the interictal period was restricted to be at least four hours away from any seizures (Fig 1). After removing segments with artifacts and segmenting the data to satisfy imposed constraints (e.g. 4-hour gap between interictal and seizure onset, etc; see Fig. 1), the amount of data for each patient corresponded to: P1: interictal, 143 hours 30 minutes, preictal 2 hours; P2: interictal 90 hours 20 minutes, preictal 2 hours; P3: interictal, 107 hours 10 minutes, 2 hours preictal; P4: interictal, 104 hours, preictal 1 hour 50 minutes; P5: interictal, 18 hours, preictal, 3 hours.

The recorded data were band-pass filtered to obtain the LFP and MUA and segmented in 4s overlapping windows, from which four groups of features were extracted (see Material and Methods and Fig 2): (i) the power spectrum of the LFP in ten defined frequency bands (LFP power spectrum), (ii) different network measures based on pairwise coherence matrix (LFP coherence), (iii) the total number of threshold crossing events (MUA count) in each window and channel, (iv) different network measures of MUA count correlation matrices (MUA correlation). The prediction task was performed with a recurrent neural network using LSTM cells [19] (see Material and Methods and Supplementary Fig 1). LSTM cells are particularly efficient for capturing long-term temporal dependencies as they do not suffer from the vanishing gradient issue [22], and have been widely used in many difficult learning problems such as speech synthesis [23], handwriting generation [24], or language translation [25]. Sequences of ordered features in time are then given as input to the LSTM for the supervised learning task.

The prediction task was performed for each patient separately. We used a leave-one-seizure-out cross-validation scheme, i.e. for each seizure, the test set was composed of the corresponding preictal and interictal periods, while the algorithm was trained on the preictal and interictal periods corresponding to all of the other seizures. (A predictive assessment based on an alternative cross-validation scheme where the sequence of labeled interictal and preictal time windows were first randomly shuffled in time, followed by the determination of cross-validation folds, i.e. training and test datasets, is shown in Supplementary Fig. 2.) For all patients, we evaluated the predictive performance by computing the AUC scores, reaching in several cases values above 0.8 (Fig. 3, AUC = 0.5 and AUC = 1 correspond to “asymptotic” chance level and perfect prediction, respectively). AUC scores were significantly better than chance, based on surrogate datasets, for most groups of features. Chance-level surrogate datasets were obtained by random permutation of the labels (interictal, preictal) assigned to each time window segment (see Materials and Methods for additional details). The corresponding chance-level AUC distributions are shown in the Supplementary Fig. 3. Prediction performance based on MUA count was on average better than chance for all patients. The best performing group of features was patient specific. An example of the prediction probabilities for the interictal and preictal segments for one test seizure in P2 is shown in Supplementary Fig 4A. While the probabilities stay high during the interictal state, an increase occurs in the preictal state. The detailed performances for each test seizure, group of features and patients are shown in Supplementary Fig 4B. We note that the hyperparameter specifying length of the history in the LSTM network was optimized directly over the test dataset, as not enough data were available to have a proper validation dataset. An assessment of the dependence of AUC scores as a function of the length of the LSTM history is shown in Supplementary Fig 5.

**Fig 3.**
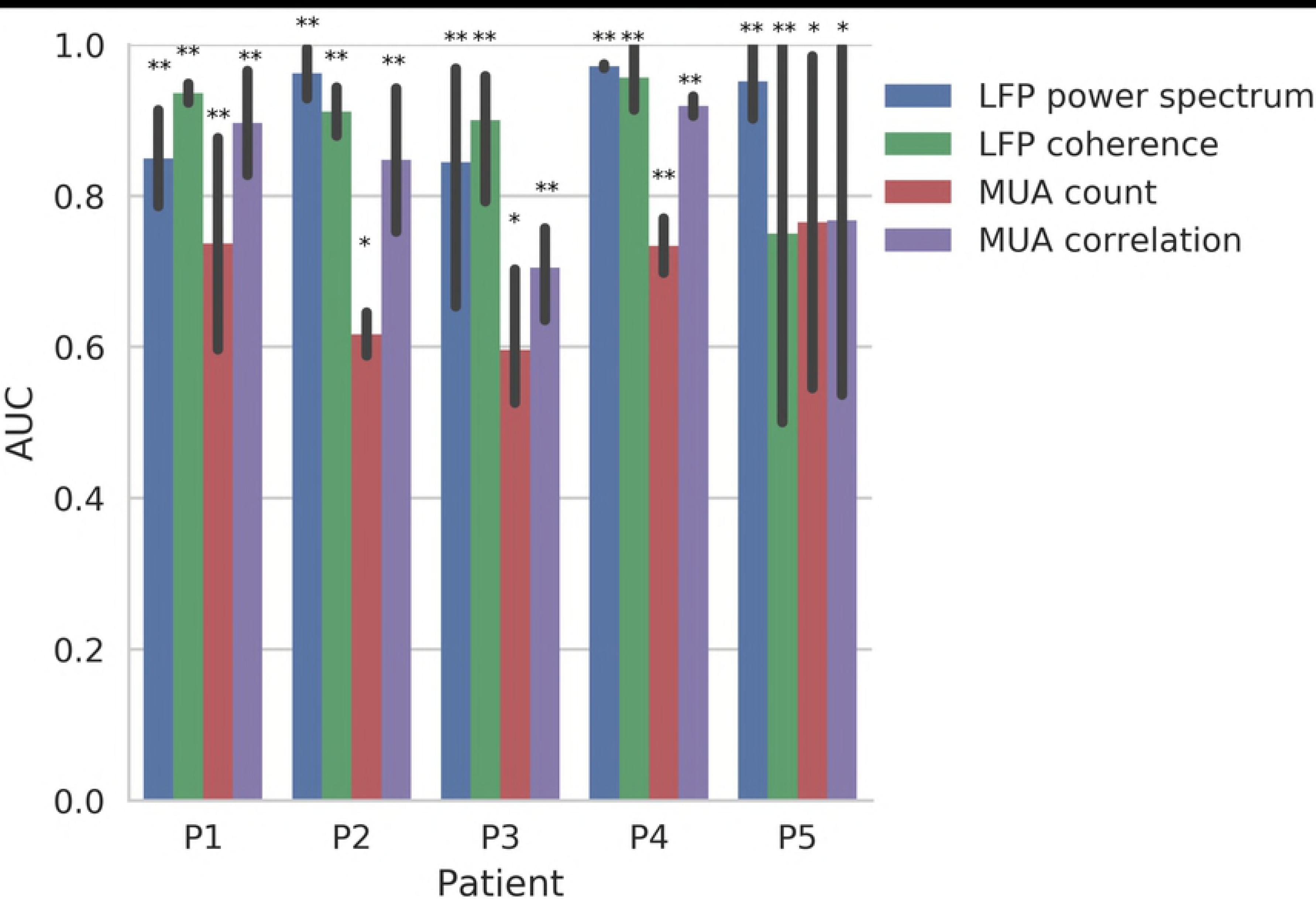
Seizure prediction performance: AUC summary including the five patients, and the four groups of features. Each bar indicates the AUC score averaged over all seizures tested in the leave-one-out cross-validation setting for each patient and each feature. The vertical lines indicate the corresponding AUC extrema. Blue: LFP power spectrum; Green: LFP pairwise spectral coherence matrix; Red: MUA count; Purple: MUA pairwise correlation matrix. Statistically significant AUCs are indicated by * for a target α = 0.05 and by ** for a target α = 0.01, after FDR multiple testing corrections (p-values were computed based on change level prediction based on surrogate data; see Materials and Methods).

**Fig 4.**
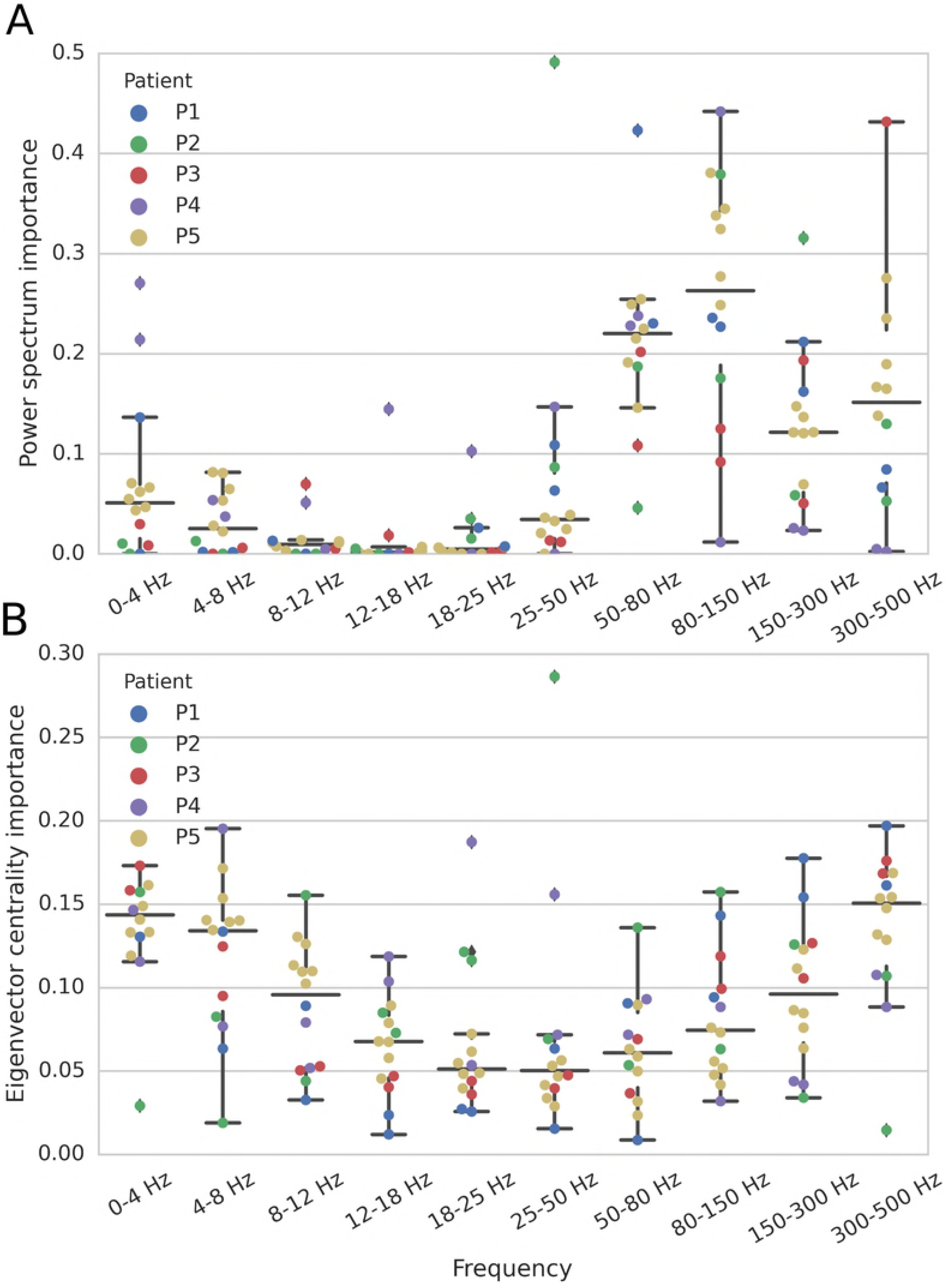
Examination of LFP feature importance by frequency band and patient. (A) Power spectrum features. (B) Features based on the leading eigenvector (eigenvector centrality) computed from the pairwise spectral coherence matrix in each frequency band. Each dot corresponds to one of the leave-one-seizure-out cross-validation folds, i.e. the interictal and preictal period of one seizure test dataset for a given patient. In these plots, center lines indicate the median, and the whiskers indicate the data point extrema. Different colors indicate different patients. Features were ranked based on the XGBoost classification algorithm.

**Fig 5.**
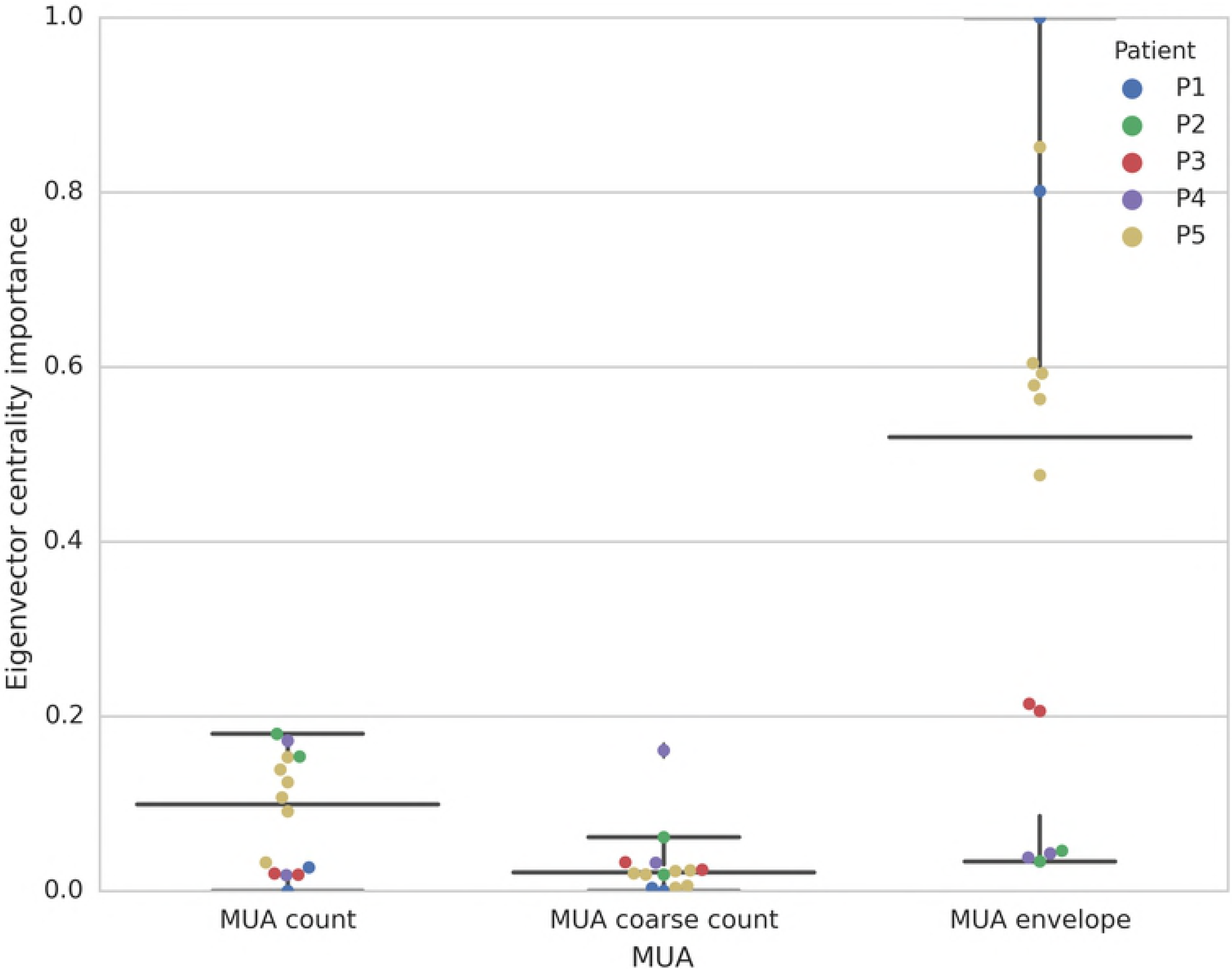
Feature importance for MUA count, MUA coarse count, and MUA envelope for all patients. Each dot corresponds to a leave one seizure-out cross-validation fold, i.e. the interictal and preictal period of one test seizure for a given patient. Center lines indicate the median, and the whiskers indicate the extreme data points. Each color corresponds to one patient.

The length of the neural recordings could extend to two weeks or more. Slow changes in the recorded signals (due to artifacts or true changes in neural activity) could occur during this period and bias the classifier, while still leading to good AUC scores, especially given the small number of seizures available. We checked that this was not the case by performing the same prediction task, but now z-scoring (zero mean, unit variance) the MUA and LFP signal for each of the original 40-minute data files. While this procedure can potentially render the classification task more difficult, as some true seizure predictive information may be lost, we were still able to obtain good AUC classification scores (Supplementary Fig. 6), confirming that the previous classifiers were not biased by slow changes in mean and variance.

**Fig 6.**
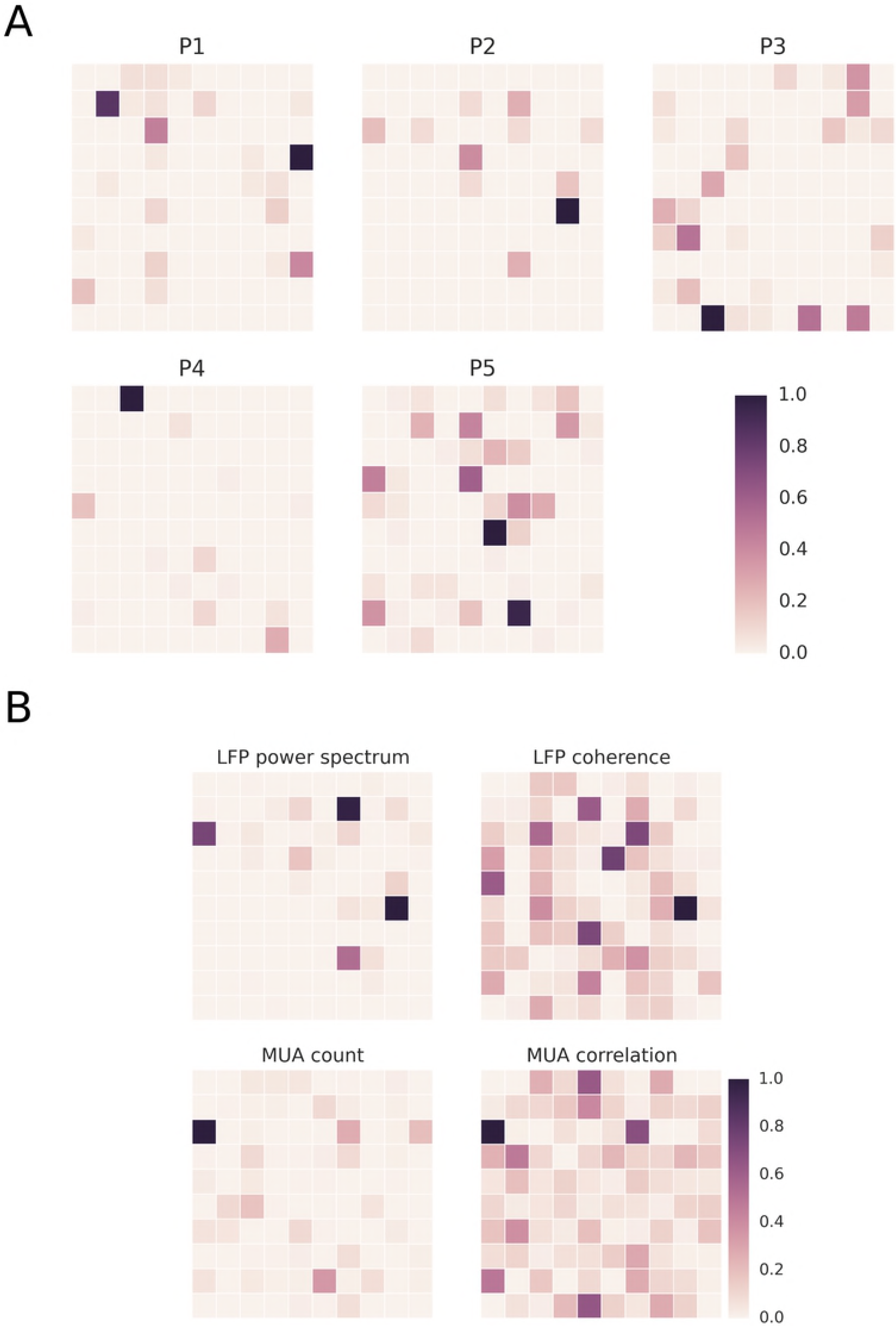
Importance of different MEA sites for seizure prediction. (A) MUA count feature. (B) Patient P3. LFP power spectrum, frequency band 300-500 Hz. The importance of a given MEA site is ranked from 0 to 1. MEA sites contributing to high importance features can be sparse and very localized in the 4 X 4 mm^2^ neocortical patch, indicating the existence of predictive microdomains of neural activity.

### Low (0-8 Hz) and high (50-500 Hz) frequency bands are the most predictive LFP features

Next, we examined if features from specific LFP frequency bands were more predictive than others. We used the XGBoost algorithm, a fast implementation of a gradient tree boosting algorithm [21], to rank the features according to their predictive power. According to this approach, the lower (0-5 Hz) and higher (50-500 Hz) frequency bands were found to be the most important features for LFP (Fig 4). This finding applied to both univariate (single channel power spectrum) and multivariate features computed from the pairwise spectral coherence matrix, i.e. the leading eigenvalue and corresponding eigenvector (eigenvector centrality). The eigenvector centrality is commonly used as an estimate of the influence of different nodes (channel) in the network as measured by the MEA.

### Most predictive MUA features subsets

As shown in Figure 3, multivariate features based on the MUA pairwise correlation matrix led to higher AUC scores in comparison to just the MUA count vector corresponding to different MEA channels or sites. These multivariate features corresponded to the leading eigenvalue and corresponding eigenvector centrality of the pairwise correlation matrices computed based on MUA count (0.5-ms time bins, 4-second time windows), coarse MUA count (100-ms time bins, 4-second time windows), and the MUA envelope (2 kHz sampling rate, 4-second time windows). As done for the LFP-based features, we also examined the importance of MUA related multivariate features based on the XGBoost algorithm. We found the leading eigenvector (eigenvector centrality) of the pairwise correlation matrix for the MUA envelope to be the more important feature in comparison to the corresponding feature in the MUA count or coarse count signals (Fig 5).

### Prediction with selected feature subsets

The same feature subsets identified with the XGBoost algorithm to be more important in average across patients, as described above, were then used as inputs to the LSTM networks for seizure prediction. The use of these selected subset of features did not improve, however, the seizure prediction performance (Supplementary Fig. 7). We note, on the other hand, that the use of this subset of these selected features reliably gave average AUC scores above 0.8 for all patients, while the best performing group of feature changed across patients in Fig 3.

### Spatial sparseness of predictive patterns

Using the same XGBoost algorithm, we also examined the spatial organization of the feature importance in the MEA, both for LFP and MUA. The algorithm singled out sparse/isolated electrodes in the MEA as the sites for the most important features, suggesting that localized patterns of activity were relevant for seizure prediction (Fig 6A). Interestingly, while univariate feature importance (LFP power spectrum, MUA count) relied on more localized patterns, the ranking of MEA spatial sites based on the eigenvector centrality computed from the LFP coherence and MUA correlation matrices was more diffuse (Fig 6B). Additionally, these ranking of MEA sites depended on the frequency band and group of features.

## Discussion

Based on localized microelectrode array recordings of MUA and/or LFPs, we have shown that human focal seizures can be predicted based on intracortical signals recorded from neocortical patches away from the identified seizure onset areas. We found that features related to low (< 8 Hz) and high (> 50 Hz) LFP frequency bands as well as multivariate features related to multi-channel MUA were the most predictive. Importantly, predictive features could be spatially sparse in the MEA suggesting that microdomains of neural activity contributed to seizure prediction.

Our previous analysis [9] showed preliminary and promising results for seizure prediction based on single-neuron spiking activity. However, the task of sorting and tracking single (neuron) units’ extracellular action potential waveforms across long periods of time is challenging. As a consequence, datasets based on single unit activity commonly span only small periods of interictal data. Here, we focused on multiunit activity, which allowed us to extend the analysis to much longer time periods. In particular, in our definition of MUA count, extracellular action potentials were detected in the usual way as done for single unit detection, except they were not sorted in different units. We also extended the analyses to new patient datasets and used a different approach for seizure prediction. In contrast to seizure prediction based on deviations from the one-class interictal state (as done in [9]), here we used supervised learning with two explicit classes, i.e. interictal and preictal states or classes of neural activity.

Our finding that neuronal network dynamics in neocortical patches distal to identified seizure onset areas are predictive of seizure initiation suggests that large-scale networks may be engaged in the neural dynamics leading to seizures. While low frequencies LFPs are thought to be generated mostly by post-synaptic potentials induced by nonlocal inputs from other regions, high frequency (> 100 Hz) LFPs are typically thought to be more local, further supporting the finding that local activity is altered [26]. Whether transient HFOs, as commonly defined in terms of narrowband oscillations, contribute to predictive features remains an issue for further investigation. The predictive features based on MUA further supported the local nature of these predictive neural dynamics. Overall, our results indicate that preictal neural activity in the recorded neocortical patches involved not only subthreshold postsynaptic potentials, perhaps driven by the distal SOAs before seizure onset, but also neuronal spiking in the local recurrent neocortical networks.

We reset the state of the LSTM between each new input to the LSTM network (i.e. an history size between 4 and 50 seconds to be consistent with the methods). It is however possible to perform continuous prediction without resetting the state of the LSTM. While no significant improvements were observed when using continuous prediction on LSTM networks on our data (results not shown), this setup is particularly well suited for real-time seizure prediction system, where the LSTM could follow slow change of the signal over long periods while retaining the critical features for seizure prediction. We also hope to provide a more systematic analysis of the correlation between spatiotemporal properties of the input features and the temporal activation of LSTM cells in a future study.

Many other machine learning applications suffer from imbalanced dataset problems, e.g. medical diagnosis [27]. In our datasets, this ratio corresponded to about 1:100. This issue was aggravated by the small sample size of our dataset, since our recordings are only two weeks long with a few seizures [28]. To overcome this problem, we used techniques such as oversampling and cost-sensitive learning. Another way to circumvent this issue would be to collect more data. Recently, long-term studies have successfully obtained intracranial ECoG recordings of periods spanning several months to years, showing the feasibility of a permanently implanted seizure forecast system, as well as provided invaluable amount of data for seizure prediction (e.g. [4]). Our results demonstrate that long-term implantation of MEA could be of interesting value as well for therapeutic purposes, and would provide the support for crowdsourcing seizure prediction algorithm, which have been proven to be highly efficient in maximizing the improvement to cost ratio of seizure prediction algorithms [7]. It would finally allow for fair comparison of seizure prediction efficacy with other modalities, i.e. ECoG and depth-electrode recordings.

Functional and diffusion MRI studies also support the view that broad and distributed network changes outside of the epileptogenic zone are associated with focal epilepsy [29]. Despite having a predictive value on surgical outcomes [30], it is still largely unknown if these functional and structural alterations contribute to triggering seizures. Computational studies have demonstrated that network alterations affect seizure onset time and areas [31,32]. Neural activity recorded by the implanted MEAs in our study might have reflected the dynamics of salience and default mode networks, which have been found to be consistently altered across patients with different epilepsy types [29]. Overall, our findings provide supporting evidence for this large-scale network view of the initiation of epileptic seizures. Our findings point also to the potential relevance of stimulation at distal brain areas, in addition to identified seizure onset areas, to improve the efficiency of closed-loop stimulation systems for seizure control and prevention. Our promising initial results warrant further studies on much larger datasets and patient groups, which we hope to conduct in the future.

## Acknowledgments

We would like to thank the patients who participated in this study and the nursing and medical staff at Massachusetts General, Brigham and Women’s, and Rhode Island Hospitals. This research was supported by: National Institute of Neurological Disorders and Stroke (NINDS), grants R01NS079533 (to WT), R01NS062092 (to SSC); the Department of Veterans Affairs, Merit Review Award I01RX000668 (to WT); and the Pablo J. Salame ‘88 Goldman Sachs endowed Assistant Professorship of Computational Neuroscience (WT). Part of this research was conducted using computational resources and services at the Center for Computation and Visualization, Brown University. The contents do not represent the views of the U.S. Department of Veterans Affairs or the United States Government.

## Supplementary Figures: Captions

**Supplementary Fig 1: LSTM cell.** The state of the cell at time *t*, *c(t)*, is updated at each time step with the former output of the cell, *h(t-1)*, and new inputs, *x(t)*, depending of the value of the input gate, *i(t).* The memory of the cell is controlled by the forget gate, *f(t)*, while the output *h(t)* is controlled by the output gate, *o(t)*.

**Supplementary Fig 2: AUC scores for the cross-validation procedure based on random permutation of the order of the time windows.** The bars indicate the AUC score averaged over all seizures for each patient and feature group according to an alternative cross-validation scheme where the order of the time windows are randomly permutated before assignment to training and test datasets. Blue: LFP power spectrum; Green: LFP pairwise spectral coherence matrix; Red: MUA count; Purple: MUA pairwise correlation matrix.

**Supplementary Fig 3: AUC chance level distributions based on surrogate datasets.** Each row corresponds to a different patient, and each column to a different group of features. The vertical line indicates the AUC score of the classifier when computed on the true (not chance-level surrogate) dataset. (See main text, Materials and Methods, for details.)

**Supplementary Fig 4: Detailed prediction scores using LFP or MUA features.** (A) Prediction probabilities corresponding to interictal/preictal periods for a single test seizure from patient P3 for the four groups of features described above. Blue: LFP power spectrum. Red: LFP pairwise spectral coherence matrix. Green: MUA count. Cyan: MUA count pairwise correlation matrix. The inset shows a zoomed-in view of the preictal prediction probabilities. (B) ROC curves based on each test seizure and each of the four group of features for the five patients. The legend indicates the average AUC scores over all seizures of a given patient for each feature group as indicated by the color.

**Supplementary Fig 5: AUC scores as a function of the length of the history.** (A) LFP power spectrum (B) LFP coherence (C) MUA count (D) MUA correlation. Each curve corresponds to the AUC average computed across the test datasets for each patient.

**Supplementary Fig 6: AUC scores for the normalized data.** The bars indicate the AUC scores averaged over all seizures tested in the leave-one-out cross-validation setting for each patient and each feature. Blue: LFP power spectrum; Green: LFP pairwise spectral coherence matrix; Red: MUA count; Purple: MUA pairwise correlation matrix.

**Supplementary Fig 7: ROC curves for all patients obtained by training the LSTM network on a subset of the most predictive features.** The legend indicates the corresponding prediction based on the average AUC score.

## References

1. England MJ, Liverman CT, Schultz AM, Strawbridge LM. Epilepsy across the spectrum: Promoting health and understanding. Epilepsy Behav. 2012;25: 266–276. doi:10.1016/j.yebeh.2012.06.016

2. Löscher W, Schmidt D. Modern antiepileptic drug development has failed to deliver: Ways out of the current dilemma: Ways Out of the Current Dilemma with New AEDs. Epilepsia. 2011;52: 657–678. doi:10.1111/j.1528-1167.2011.03024.x

3. Engel J. A greater role for surgical treatment of epilepsy: why and when? Epilepsy Curr. 2003;3: 37–40.

4. Cook MJ, O’Brien TJ, Berkovic SF, Murphy M, Morokoff A, Fabinyi G, et al. Prediction of seizure likelihood with a long-term, implanted seizure advisory system in patients with drug-resistant epilepsy: a first-in-man study. Lancet Neurol. 2013;12: 563–571. doi:10.1016/S1474-4422(13)70075-9

5. Heck CN, King-Stephens D, Massey AD, Nair DR, Jobst BC, Barkley GL, et al. Two-year seizure reduction in adults with medically intractable partial onset epilepsy treated with responsive neurostimulation: Final results of the RNS System Pivotal trial. Epilepsia. 2014;55: 432–441. doi:10.1111/epi.12534

6. Morrell MJ. Responsive cortical stimulation for the treatment of medically intractable partial epilepsy. Neurology. 2011;77: 1295–1304. doi:10.1212/WNL.0b013e3182302056

7. Kuhlmann L. Crowdsourced epileptic seizure prediction with big data. 2017;

8. Schevon CA, Weiss SA, McKhann G, Goodman RR, Yuste R, Emerson RG, et al. Evidence of an inhibitory restraint of seizure activity in humans. Nat Commun. 2012;3:1060. doi:10.1038/ncomms2056

9. Truccolo W, Donoghue JA, Hochberg LR, Eskandar EN, Madsen JR, Anderson WS, et al. Single-neuron dynamics in human focal epilepsy. Nat Neurosci. 2011;14: 635–641. doi:10.1038/nn.2782

10. Truccolo W, Ahmed OJ, Harrison MT, Eskandar EN, Cosgrove GR, Madsen JR, et al. Neuronal Ensemble Synchrony during Human Focal Seizures. J Neurosci. 2014;34: 9927–9944. doi:10.1523/JNEUROSCI.4567-13.2014

11. Wagner FB, Eskandar EN, Cosgrove GR, Madsen JR, Blum AS, Potter NS, et al. Microscale spatiotemporal dynamics during neocortical propagation of human focal seizures. NeuroImage. 2015;122: 114–130. doi:10.1016/j.neuroimage.2015.08.019

12. Bartolomei F, Lagarde S, Wendling F, McGonigal A, Jirsa V, Guye M, et al. Defining epileptogenic networks: Contribution of SEEG and signal analysis. Epilepsia. 2017;58: 1131–1147. doi:10.1111/epi.13791

13. Kramer MA, Cash SS. Epilepsy as a disorder of cortical network organization. The Neuroscientist. 2012;18: 360–372.

14. Spencer SS. Neural Networks in Human Epilepsy: Evidence of and Implications for Treatment. Epilepsia. 2002;43: 219–227.

15. Stark E, Abeles M. Predicting Movement from Multiunit Activity. J Neurosci. 2007;27: 8387–8394. doi:10.1523/JNEUROSCI.1321-07.2007

16. Mitra PP, Pesaran B. Analysis of dynamic brain imaging data. Biophys J. 1999;76: 691–708.

17. Brinkmann BH, Wagenaar J, Abbot D, Adkins P, Bosshard SC, Chen M, et al. Crowdsourcing reproducible seizure forecasting in human and canine epilepsy. Brain. 2016;139: 1713–1722. doi:10.1093/brain/aww045

18. Gers FA, Schmidhuber J, Cummins F. Learning to forget: continual prediction with LSTM. Neural Comput. 12: 2451–2471.

19. Hochreiter S, Schmidhuber J. Long Short-Term Memory. Neural Comput. 1997;9: 1735–1780. doi:10.1162/neco.1997.9.8.1735

20. Benjamini Y, Hochberg Y. Controlling the false discovery rate: a practical and powerful approach to multiple testing. J R Stat Soc. 1995;57: 289–300.

21. Chen T, Guestrin C. Xgboost: A scalable tree boosting system. Proceedings of the 22Nd ACM SIGKDD International Conference on Knowledge Discovery and Data Mining. ACM; 2016. pp. 785–794. Available: http://dl.acm.org/citation.cfm?id=2939785

22. Bengio Y, Simard P, Frasconi P. Learning Long-Term Dependencies with Gradient Descent is Difficult. IEEE Trans Neural Netw. 1994;5: 157–166.

23. Fan Y, Qian Y, Xie F-L, Soong FK. TTS synthesis with bidirectional LSTM based recurrent neural networks. Interspeech. 2014. pp. 1964–1968. Available: https://mazsola.iit.uni-miskolc.hu/~czap/letoltes/IS14/IS2014/PDF/AUTHOR/IS140552.PDF

24. Graves A, Mohamed A, Hinton G. Speech recognition with deep recurrent neural networks. Acoustics, speech and signal processing (icassp), 2013 ieee international conference on. IEEE; 2013. pp. 6645–6649. Available: http://ieeexplore.ieee.org/abstract/document/6638947/

25. Luong M-T, Sutskever I, Le QV, Vinyals O, Zaremba W. Addressing the rare word problem in neural machine translation. ArXiv Prepr ArXiv14108206. 2014; Available: https://arxiv.org/abs/1410.8206

26. Jiruska P, Alvarado-Rojas C, Schevon CA, Staba R, Stacey W, Wendling F, et al. Update on the mechanisms and roles of high-frequency oscillations in seizures and epileptic disorders. Epilepsia. 2017; doi:10.1111/epi.13830

27. Lichman M. UCI Machine Learning Repository. Univ Calif Irvine Sch Inf Comput Sci. 2013;

28. Japkowicz N, Stephen S. The Class Imbalance Problem: A Systematic Study. Intell Data Anal. 2002;6: 429–50.

29. Besson P, Bandt KS, Proix T, Lagarde S, Jirsa VK, Ranjeva J-P, et al. Anatomic consistencies across epilepsies: a stereotactic-EEG informed high-resolution structural connectivity study. Brain. 2017;

30. Keller SS, Glenn GR, Weber B, Kreilkamp BAK, Jensen JH, Helpern JA, et al. Preoperative automated fibre quantification predicts postoperative seizure outcome in temporal lobe epilepsy. Brain. 2017;140: 68–82. doi:10.1093/brain/aww280

31. Petkov G, Goodfellow M, Richardson MP, Terry JR. A Critical Role for Network Structure in Seizure Onset: A Computational Modeling Approach. Front Neurol. 2014;5. doi:10.3389/fneur.2014.00261

32. Sinha N, Dauwels J, Kaiser M, Cash SS, Brandon Westover M, Wang Y, et al. Predicting neurosurgical outcomes in focal epilepsy patients using computational modelling. Brain. 2017;140: 319–332. doi:10.1093/brain/aww299

